# Ecological and evolutionary adaptations shape the gut microbiome of BaAka African rainforest hunter-gatherers

**DOI:** 10.1101/019232

**Authors:** Andres Gomez, Klara Petrzelkova, Carl J. Yeoman, Michael B. Burns, Katherine R. Amato, Klara Vlckova, David Modry, Angelique Todd, Carolyn A. Jost Robinson, Melissa J. Remis, Manolito Torralba, Karen E. Nelson, Franck Carbonero, H. Rex Gaskins, Brenda Wilson, Rebecca M. Stumpf, Bryan A. White, Steven R. Leigh, Ran Blekhman

**Affiliations:** Department of Genetics, Cell Biology, and Development, University of Minnesota, Twin Cities, MN, 55108; Department of Ecology, Evolution and Behavior, University of Minnesota, Twin Cities, MN, 55108; Institute of Vertebrate Biology, Academy of Sciences of the Czech Republic, Brno, Czech Republic; Department of Pathology and Parasitology, Faculty of Veterinary Medicine, University of Veterinary and Pharmaceutical Sciences, Brno, Czech Republic; Institute of Parasitology, Biology Center of the Academy of Sciences of the Czech Republic, Českè Budějovice, Czech Republic; Liberec Zoo, Liberec, Czech Republic; Department of Animal and Range Sciences, Montana State University, Bozeman, MT, 59717; Department of Anthropology, University of Colorado, Boulder, CO, 80309, USA; CEITEC, Central European Institute for Technology, Brno, Czech Republic; World Wildlife Fund, Dzanga-Sangha Protected Areas, Bayanga, Central African Republic; Department of Anthropology, University of North Carolina, Wilmington, NC, 28403; Deptartment of Anthropology, Purdue University, West Lafayette, IN, 47907; The J. Craig Venter Institute, Rockville, MD, 20850; Institute for Genomic Biology, University of Illinois at Urbana Champaign, IL, 61801; Department of Animal Sciences, University of Illinois at Urbana Champaign, IL, 61801; Department of Food Science, University of Arkansas, Fayetteville, AK, 72704; Department of Microbiology, University of Illinois at Urbana-Champaign, IL, 61801; Department of Anthropology, University of Illinois at Urbana Champaign, IL, 61801

**Keywords:** BIOLOGICAL SCIENCES, Anthropology, Ecology, Evolution, Microbiology, BaAka hunter-gatherers, Pygmy, gut microbiome, adaptation, evolution

## Abstract

The gut microbiome provides access to otherwise unavailable metabolic and immune functions, likely affecting mammalian fitness and evolution. To investigate how this microbial ecosystem impacts evolutionary adaptation of humans to particular habitats, we explore the gut microbiome and metabolome of the BaAka rainforest hunter-gatherers from Central Africa. The data demonstrate that the BaAka harbor a colonic ecosystem dominated by Prevotellaceae and other taxa likely related to an increased capacity to metabolize plant structural polysaccharides, phenolics, and lipids. A comparative analysis shows that the BaAka gut microbiome shares similar patterns with that of the Hadza, another hunter-gatherer population from Tanzania. Nevertheless, the BaAka harbor significantly higher bacterial diversity and pathogen load compared to the Hadza, as well as other Western populations. We show that the traits unique to the BaAka microbiome and metabolome likely reflect adaptations to hunter-gatherer lifestyles and particular subsistence patterns. We hypothesize that the observed increase in microbial diversity and potential pathogenicity in the BaAka microbiome has been facilitated by evolutionary adaptations in immunity genes, resulting in a more tolerant immune system.

**Significance:** Human ecological adaptation requires changes at the genomic level. However, the gut microbiome, the collection of microbes inhabiting the gastrointestinal tract and their functions, also responds significantly to ecological challenge. To determine how the gut microbiome responds to evolutionary adaptations in the host, we profiled gut bacterial communities of the BaAka, rainforest hunter-gatherers from Central Africa. The gut microbiome of the BaAka shows adaptations to metabolize foods rich in fiber, tannins and fats. Similarly, higher bacterial diversity and abundance of pathogenic bacteria, compared to other hunter-gatherers and western populations, suggest that the BaAka immune system evolved to coexist with increased pathogen threats. Accordingly, these results show how the gut microbiome contributes to human ecological plasticity, impacting host adaptation and evolution.

---

Particular gut microbiome patterns have been linked to specific physiological phenotypes in humans (1, 2). Recently, the characterization of gut microbiomes across geographically and culturally diverse human populations have helped researchers pinpoint the effects of lifestyle factors on the composition of these microbial communities and the potential influence of gut microbes in human health and disease (3-5). For instance, specific gut microbiome assemblages, likely supported by westernized diets and industrialized lifestyles, are believed to play a role in the high incidence of metabolic disorders observed in some modern human populations (6). These patterns are opposed to the more ancestral microbiome traits seen in traditional societies (4). Thus, comparative analyses of diet-microbiome interactions facilitate an understanding of diverse lifestyles and diets on human adaptation and health.

With respect to population-specific adaptations, persistent debate has accompanied the question of whether or not forest foraging lifestyles among Central African rainforest hunter-gatherer populations supply adequate nutrition, in the context of specific phenotypes and ecological challenges (7-9). In this regard, small human body size, a characteristic common to some rainforest hunter-gatherer populations (the “pygmy” phenotype), has been variably hypothesized to have evolved in response to energetic limitations, forest density, high heat and humidity, increased parasite load and high adult mortality (10-12). Given the role of the gut microbiome in harvesting nutrients from low energy foods and their critical impact in immune balance, understanding the gut microbiome of these hunter-gatherer populations may provide insights into their ecology and evolutionary adaptations.

Here, we explore gut microbiome and metabolome composition in stool samples of the BaAka, forest foragers from the Dzanga Sangha Protected Areas (DSPA), Central African Republic. A comparative analysis of microbiome composition was conducted, including the Hadza hunter-gatherers from Tanzania, urban living Italians (13) and US Americans (2). Thus, we predict that the characterization bacterial communities and metabolomes in the BaAka gut contributes to persistently debated anatomical, genetic, and physiological specializations that help distinguish forager populations from non-traditional western societies.

## Results

### The BaAka gut microbiome

Gut microbiome composition was profiled through 454-16S rRNA pyrosequencing of stool samples from 27 BaAka. After quality control, we obtained a total of 151,407 pyrotag reads, with an average number of 5,607 ± 1,841 reads per sample (range 2,939-10,673). Firmicutes and Bacteroidetes dominated the BaAka gut microbiome (45 ± 16.9 and 39 ± 20.6 % respectively) (Figure 1a). Unclassified bacteria comprised 10.8 ± 14.4% of all reads, whereas Proteobacteria and Spirochaetes were the most abundant among minor phyla (2.3 ± 2.9 and 1.6 ± 2.2 % respectively).

**Figure 1.**
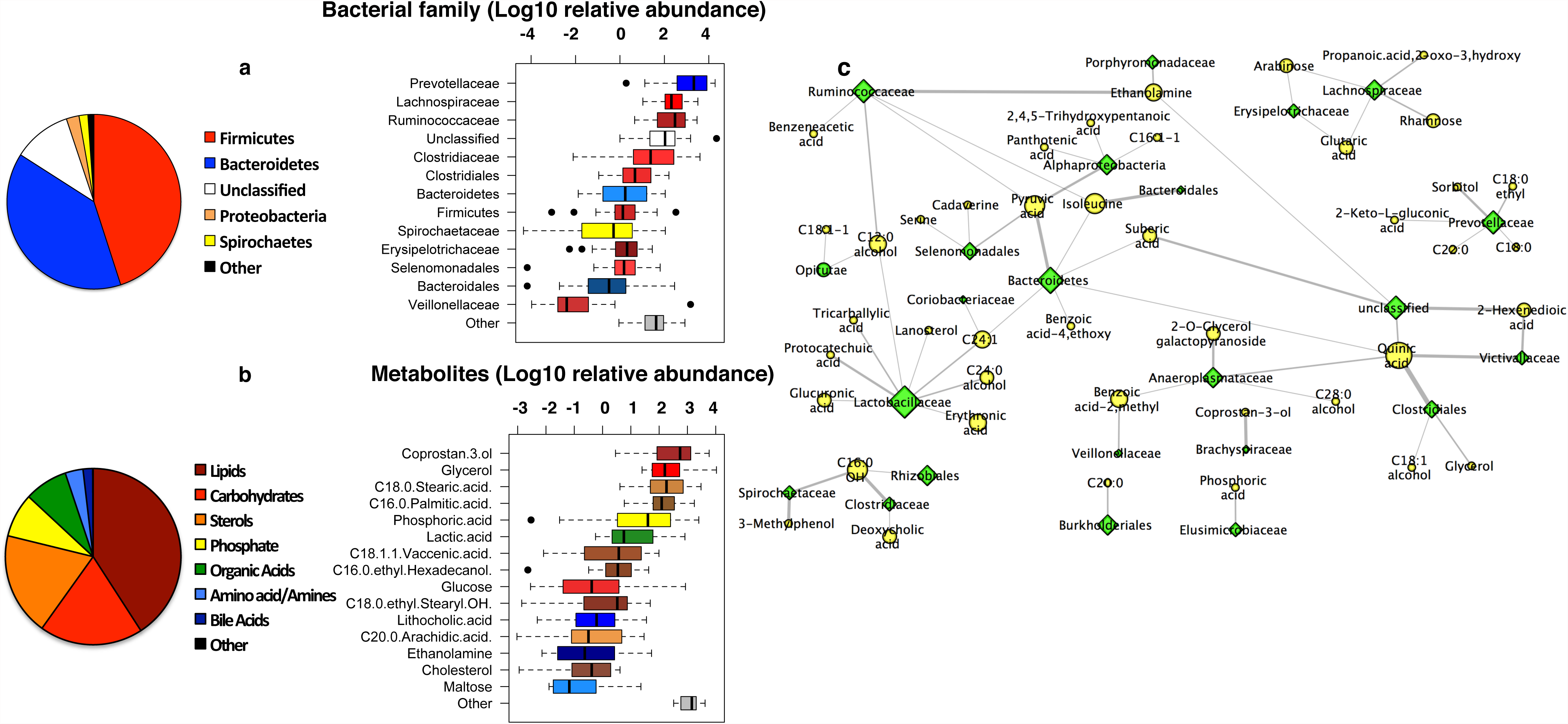
Gut microbiome and metabolome composition in the BaAka. (**a**)Relative abundance of taxa at phyla and family level. (**b**)Relative abundance of metabolites according to broad and specific classifications. (**c**) Subnetwork view of gut microbiome-metabolome interactions in the BaAka. Green and yellow nodes represent bacterial taxa (at the family level) and metabolites respectively. Edges represent Spearman correlation coeficients between metabolites and families (*rho* > 0.5). Node size is proportional to the number of connections, and edge width represents strength of the correlation.

A taxonomic analysis at family level detected 55 taxa in the BaAka microbiome, with Prevotellaceae (mainly *Prevotella* spp.) identified as the predominant family (33.6 ± 23.2 %) (Figure 1a). High abundances of Prevotellaceae (up to 53% in average) are also predominant in other African populations that rely on high-fiber content diets (14, 15). Together, members of the Clostridiales order (Lachnospiraceae (13.31 ± 8.49%), Ruminococcaceae (13 ± 8%), Clostridiaceae (8.43 ± 9.35%), and unclassified Clostridiales (2.79 ± 2.27%)) comprised the second most abundant taxa (37.5 ± 28.6 %), while other members of the Bacteroidetes (mainly unclassified Bacteroidales) reached 3.51 ± 4.5 % relative abundance. Spirochaetaceae (primarily *Treponema* spp.) was the most abundant among minor taxa comprising 1.6 ± 2.2 % in the BaAka.

### The BaAka gut metabolome and microbiome-metabolome interactions

From the 27 samples analyzed, 20 yielded usable gas chromatography/mass spectrometry (GC/MS) spectra. Analysis of both polar and non-polar metabolites in these stool samples detected a total of ∼2,500 mass spectral features, and from these, 260 compounds were positively identified. After eliminating metabolites found in less than 50% of samples, the stool metabolomes of the BaAka were mainly grouped into lipids (40.8%), carbohydrates (19%), sterols (18.9%), phosphates (8%), organic acids (7.9%), amino acids and amines (3.3%), and bile acids (1.7%) (Figure 1b). Coprostan-3-ol, the main conversion product from cholesterol metabolism in the distal colon (16), was the most abundant metabolite (15.6 ± 10.6%), followed by glycerol (12.7 ± 11.74%), stearic acid (11.9 ± 8.7%), palmitic acid (10 ± 6.5%), phosphoric acid (8 ± 8.5%) and lactic acid (4.5 ± 5%). Other metabolites comprised 2% of abundance or less (Figure 1b).

Spearman correlations between metabolites (relative abundance) and taxa (at family level) exhibiting *rho*≥0.5 (false discovery rate, q<0.05) (**Table S1**) were chosen to visualize microbiome-metabolite interactions and infer potential microbiome functions (Figure 1c). The most dominant taxa in the BaAka gut established correlations with abundances of metabolites involved in plant cell wall processing and phenolics. For instance, Prevotellaceae formed the strongest associations with sorbitol, a sugar alcohol and a product of glucose reduction and with keto-L-gluconic acid, another product of glucose metabolism. Abundances of Lachnospiraceae correlated with arabinose and rhamnose, structural components of pectic polysaccharides and hemicelluloses, and with propanoic acid, a product of polysaccharide fermentation. Ruminococcaceae showed strong associations with ethanolamine and pyruvic acid, both metabolites derived from energetic turnover mechanisms, and with benzeneacetic acid, an aromatic compound likely associated to the phenolic fraction of plants (17). The same is the case for Spirochaetaceae, which formed associations with methyl-phenol. In this regard, quinic acid, also a plant polyphenolic, formed several strong connections with unclassified Clostridiales and Bacteroidetes, and also with Victivalllaceae, and Anaeroplasmataceae, both of which have been related to increased plant matter degradation in the herbivore gut (18, 19). Other abundant taxa in the BaAka such as Clostridiaceae and Lactobacillaceae also formed strong associations with palmitic acid, deoxycholic acid (a secondary bile acid) and with other metabolites associated with lipid processing (Figure 1c).

### The BaAka gut microbiome in comparison to that of other hunter-gatherer and non-western societies

To understand the composition of the BaAka gut microbiome in the context of other populations and diets, we incorporated data from another hunter-gatherer group, the Hadza from Tanzania, and two western populations, Italians and US Americans. These data indicate that the BaAka gut microbiome exhibits significantly higher richness and diversity (P<0.001, Wilcoxon rank-sum test) (Figure 2a) and less inter-individual variation than those of the Hadza hunter-gatherers from Tanzania, Italians, and Americans from the US (Figure 2b; Permutation test for difference in medians, P<0.001).

**Figure 2.**
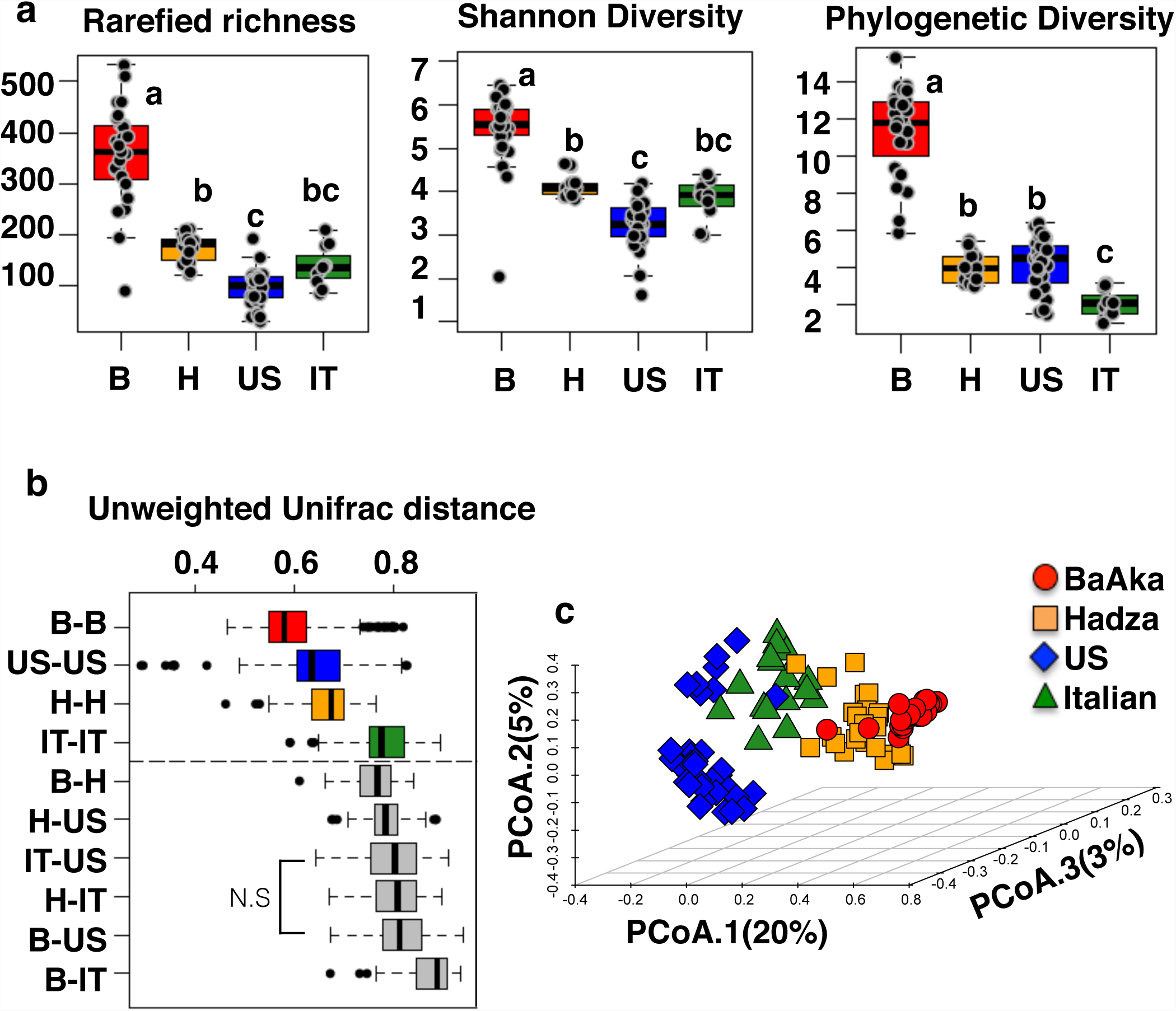
Comparison of the BaAka gut microbiome against that of the Hadza, USA americans, and italians. **(a)** Diversity indexes; rarefied richness, shannon diversity and phylogenetic diversity (1,000 reads per sample). Different letters denote significant differences (*P*<0.05) according to Kruskal-Wallis (adjusted) pair-wise comparisons. **(b)** Pair-wise unweighted unifrac distances within (upper panel) and between groups (lower panel). All within and between pairwise distances are significant (Differneces in medians, permutation test P<0.001) unless stated in the grapgh (N.S). **(c)** Principal coordinates analysis based on unweighted unifrac distances. Each symbol corresponds to the 16S rRNA bacterial community composition in one stool sample.

Although the gut microbiomes of the BaAka were more similar to those of the Hadza than they were to those of the western groups (Figures 2b, 2c **and S1**), some taxa were significantly more abundant only in the BaAka (FDR-adjusted Kruskal-Wallis tests, q<0.01). This is the case of Clostridiaceae, Anaeroplasmataceae, and Victivallaceae (Figures 3, **S2a, S2b, and table S2**). Ribosomal sequences of these taxa were all related to bacteria with increased capacity for metabolizing plant glycosides (18-20). Unclassified sequences affiliated to the Verrucomicrobia and Alphaproteobacteria; also increased only in the BaAka, have been previously reported as abundant in stool samples of mountain gorillas, which rely heavily on high fiber diets (21), and in the gut microbiome of children from Bangladesh, whose diet is high in plant polysaccharides (22) (**Figure S2 and table S1**).

**Figure 3.**
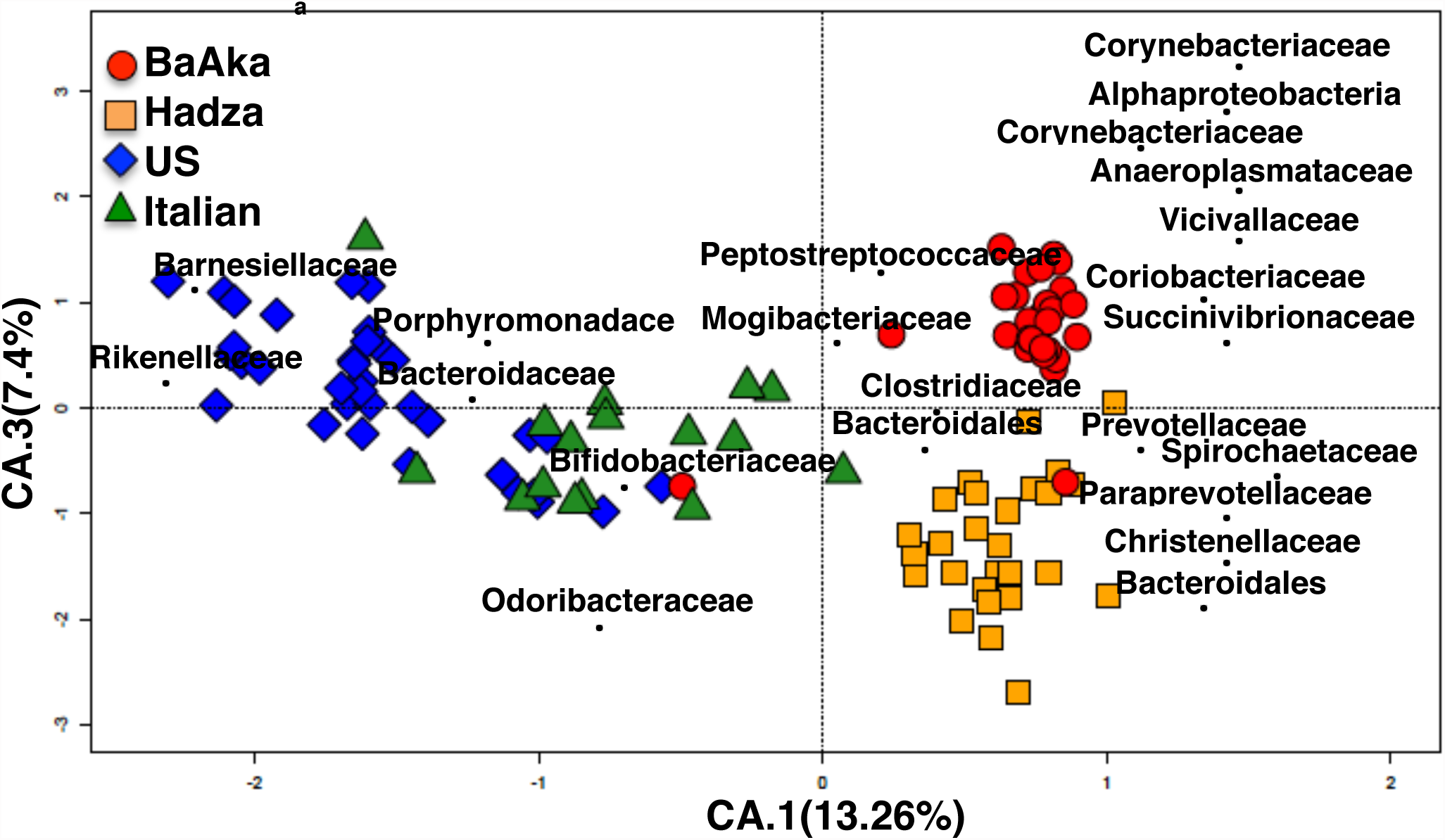
Taxa (at family level), driving gut microbiome composition differences among BaAka, Hadza, Americans (HMP) and Italians. Correspondence analysis (CA) showing indicator taxa in each group, the short distance between each taxa point and symbols gives an estimate of the relative abundance of such taxa in a given sample. For this analysis only significant discriminators, based on indicartor taxa analysis were used (see Methods). Axes 1 and 3, explaining 13.2 and 7.4% of the variation are shown separating the 4 groups. Axis 2 (11.4%) does not show a separation between BaAka and Hadza (**Figure S4**). The abundance and significance of all indicator taxa within a given group was corroborated using q-values (**Table S1**) and visulaized in boxplots. (**Figure S3**).

Other families such as Mogibacteriaceae, Corynebacteriaceae, Coriobacteriaceae, and Peptostreptococcaceae were also significantly more abundant in the BaAka only. Although these taxa can be commonly found in the mammalian gut, they all have also been associated to potential inflammation or infection (23-26). Taxa related to Odoribacteraceae were completely absent in the BaAka compared to all other groups.

Among the taxa that were significantly more enriched in both African groups (BaAka and Hadza) than in westerners (FDR-adjusted Kruskal-Wallis tests, q<0.01) (Figures 3, **S2 and Table S1**) we found Prevotellaceae, Paraprevotellaceae, Spirochaetaceae, unclassified Bacteroidales, and Succinivibrionaceae. These taxa appear to be consistently enriched in other African foragers and farmers, as well in other non-western traditional societies (3, 5, 13, 14).

Bacteroidaceae (mainly *Bacteroides*), Porphyromonadaceae, Barnesiellaceae and Rikenellaceae were more abundant in Americans and Italians (FDR-adjusted Kruskal-Wallis tests, q<0.01) compared to the African groups (Figures 3, **S2 and Table S1**). These closely related taxa (27) are believed to be signatures of western gut microbiomes (28, 29). Bifidobacteriaceae was significantly more enriched in Italians and almost absent in all other groups (FDR-adjusted Kruskal-Wallis tests, q<0.01) (Figures 3, **S2 and Table S1**).

### Microbiome functional comparison across populations

To predict the potential functional roles played by gut bacterial communities in the BaAka compared to all other groups and in both African hunter-gatherers *vs.* westerners, we used the PICRUSt (Phylogenetic investigation of communities by reconstruction of unobserved states) open source software (30). Increased predicted abundance of genes involved in infection pathways (*Staphylococcus aureus* and *Vibrio cholerae*) distinguished the BaAka over all other groups (FDR-adjusted Kruskal-Wallis tests, q<0.01) (Figure 4). Functional predictions also revealed that the gut microbiome of both western groups were significantly more enriched in genes involved in carbohydrate metabolism; galactose, starch and sucrose and the pentose phosphate pathway compared with the Hadza and the BaAka (FDR-adjusted Kruskal-Wallis tests, q<0.01).

**Figure 4.**
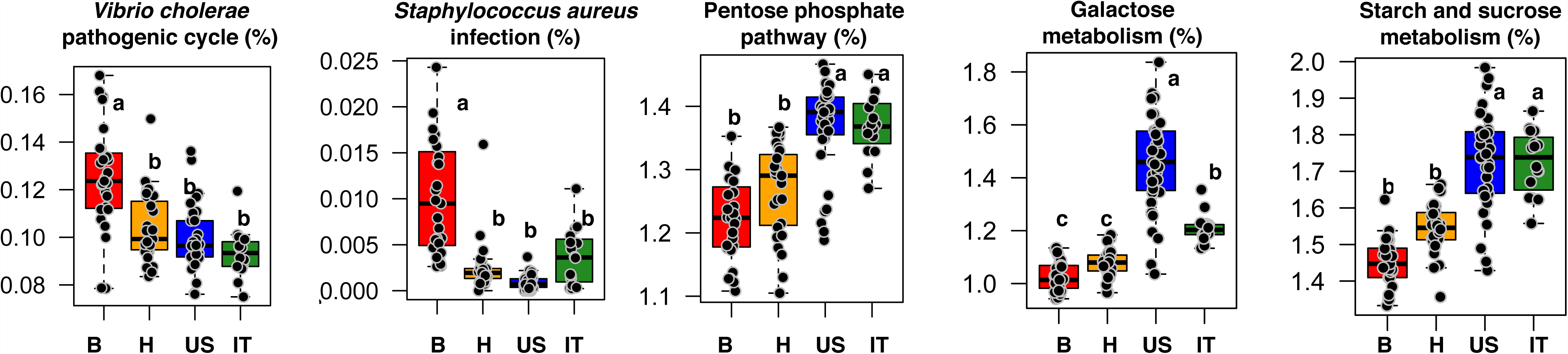
Predicted functional potential characterizing the gut microbiome of the BaAka, African and western groups. Functional predictions were made with PICRUSt. Different letters denote significant differences (*P*<0.05) according to Kruskal-Wallis (adjusted) pair-wise comparisons. B=BaAka pygmies, H=Hadza, IT=Italians, US=US Americans.

## Discussion

The present data indicate that the intestinal microbiome of the BaAka foragers may harbor compositional and functional features that likely reflect their subsistence, lifestyle, and ecology. In addition, through comparisons with the Hadza hunter-gatherers, US Americans and Italians, the data show microbiome patterns that distinguish traditional foragers from western societies. Three main features seem to characterize the gut microbiome of the BaAka hunter-gatherers, an increased capacity for processing structural polysaccharides and phenolics, a metabolome dominated by lipid-derived markers and high gut bacterial diversity, including prevalence of potential pathogens.

### The BaAka gut microbiome shows adaptations for processing phenolic-rich fibrous foods

The observation of a colonic ecosystem adapted to break down fiber may be related to the important role of fibrous polysaccharides such as bitter manioc root, yams, leaves (*Gnetum Africanum*) and wild fruits in the BaAka diet (31). Starches and wild leaves and fruits (rich in fiber and phenolics (32)) seem to be staple foods for hunter-gatherers in tropical rainforests (33, 34) and for the BaAka in Central African Republic and southeast Congo (31, 35).

Along these lines, the dominance of Prevotellaceae in the BaAka gut may be associated with large influx of xylane, pectin, and starch into their colonic ecosystem (36, 37). High abundances of Prevotellaceae, which were also shared with the Hadza, have also been observed in the gut microbiome of other rural farming groups in Africa and South America, who depend on fibrous starches for subsistence (3-5, 13-15). Prevalence of Clostridiaceae, Ruminococcaceae and Lachnospiraceae and the correlations seen between these taxa and plant polyphenolics such as benzenes, protocatechuic acid, quinic acid and other metabolites derived from fiber processing, might further support an increased capacity to break down phenolic-rich fibers in the BaAka (38).

The shared patterns seen between the BaAka and Hadza, in comparison to those in US Americans and Italians demonstrate a distinctive non-western gut microbiome. This observation is supported by dominance of other potentially fibrolytic taxa such as Succinivibrionaceae (39) and Spirocahetaceae (who was also significantly associated to abundances of phenolics) in the 2 hunter-gatherer groups. Interestingly, prevalence of Spirochaetaceae has not only been seen in the gut microbiome of other rural and traditional societies worldwide (3-5, 13, 22), but also, along with Prevotellaceae, in coprolites from extinct humans in northern Mexico (1,400 years B.P) (40).

Thus, Prevotellaceae and Spirochaetaceae may represent more ancestral gut microbiome states, specifically adapted to fiber processing. Moreover, these two taxa have also been consistently found in lowland and mountain gorillas as well as in chimpanzees (21, 41, 42), humans’ closest evolutionary relatives. The fact that these taxa are notably low in abundance or absent in westerners (4, 5), implies that large-scale agriculture and industrialization has caused loss of fiber processing diversity in modern microbiomes in favor of taxa likely specialized in metabolizing more refined carbohydrates. This is supported by the higher abundance of predicted carbohydrate processing metabolic pathways (starch, sucrose and galactose) and other closely related taxa from the Bacteroidales such as Barnesiellaceae, Porphyromonadaceae and Rikenellaceae in Italians and US Americans.

### The BaAka metabolome shows high lipid processing activity

It is notable that fatty acids and metabolites derived from lipid processing were the predominant metabolites in the BaAka. Thus, the BaAka may incorporate significant amounts fat in their diet, probably from wild meat, fish or lipid-rich nuts. Indeed, nuts from *Irvingia wombolu* constitute one of the main staple foods for the BaAka (31). These foods are richer in polyunsaturated and monounsaturated fats, compared to the predominantly saturated fats found in raised cattle (43). The combination of these unsaturated fats along with plant foods rich in fiber and polyphenolics (reported to have hypolipidemic and antioxidant effects (44)) are believed to be associated with the lower incidence of western-like metabolic diseases in hunter-gatherers (45). Furthermore, consistent with the microbiome-metabolome interaction network presented herein, colonic bacteria also interact with fatty acids to mediate conversion of other metabolites with potential antioxidant properties, such as conjugated linoleic acid (C18:2) and stearic acid (C18:0) (46), the most abundant fatty acids in the BaAka metabolome.

These metabolome-microbiome markers also motivate questions regarding energy storage and expenditure patterns in the BaAka. Metabolomes high in long-chain fatty acids and cholesterol derived markers as well as interactions of these metabolites with abundances of Lacobacillaceae have also been observed in mice and humans under high fat and high-energy diets (47). Thus, although energy intake has been suggested to be low in the wider BaAka community (35), which may result from limited food access (10, 48), these data indicate that the BaAka may not be energetically deficient and that their gut microbiomes may contribute significantly to energy harvest. However, it is unclear whether these patterns are exclusive of this group of individuals or if these observations are applicable to a wider BaAka population in light of strong fluctuations in resource availability they experience (31).

### The BaAka gut microbiome shows increased diversity and prevalence of potentially pathogenic bacteria

Even though the gut microbiome of the BaAka shares metabolic adaptations to increased fiber processing with other non-western traditional groups, these foragers show significantly higher bacterial diversity and abundance of potential pathogens compared with the Hadza, Italians and US Americans. Increased bacterial diversity in the colonic ecosystem has generally been related to dietary diversity and higher fiber intakes (49). However, the Hadza, who show significantly lower bacterial diversity compared to the BaAka, also exhibit these dietary patterns (13). Thus, another possible explanation for the observed increase in bacterial diversity in the BaAka over all other groups could be related to immunological benefits and adaptive tools to counteract potential pathogen threats (50). In this regard, increased parasite loads and infectious diseases are more common in populations inhabiting tropical rain forest (51). This may be the case of the BaAka (10, 12), who could benefit from the completive exclusion mechanisms offered by a rich bacterial ecosystem. In deed, long-term monitoring of gastrointestinal parasites in these BaAka revealed that most of them suffer from multiple infections of pathogenic gastrointestinal parasites, *i.e. Trichuris trichuira*, *Necator* spp, *Ascaris lumbricoides*, *Strongyloides* spp., in many cases in high intensities (Petrzelkova at al., unpublished data). Nevertheless, it remains to be proven whether the savanna-woodland habitats occupied by the Hadza, which make them less susceptible to gastrointestinal pathogens (52) also make them less dependent on higher bacterial diversity to neutralize pathogens compared with the BaAka.

Notably, genetic variation data indicate evolutionary adaptations in genes causing negative regulation of cytokine signaling and hence, potentially increased susceptibility to infection in the BaAka (53). Such data have not ben reported for the Hadza. This phenomenon may favor a more tolerant mucosal immune system, which may correlate with the observed increased microbiome richness, pathogen prevalence and predicted abundance of bacterial infection genes. Thus, further research should focus on whether high abundance of potential pathogens in the BaAka and a highly diverse gut microbiome are related to long-term, co-evolutionary host-microbe interactions, favored by more tolerant immune mechanisms (54).

That the gut microbiomes of the BaAka were significantly less variable than those of all other groups may also reflect strong selection pressures shaping their gut bacterial communities. Although still controversial, natural selection in West African foragers like the BaAka is believed to have favored genomic signatures involved in growth hormones (55), shorter life span and immunity (53). These may be important factors behind their small body size and high pathogen loads (10, 48). If these traits evolved in response to the ecological challenges faced by the BaAka in a dense, humid, resource-limited, and pathogen-prone tropical forest, it is likely that low inter-individual variability in gut microbiomes, high bacterial diversity and the microbiome-metabolome traits presented here are also adaptations providing the BaAka with key nutritional and immunological adaptive tools.

### Concluding remarks and future perspectives

Besides microbiome and metabolomes adaptations to particular ecological and lifestyle patterns in the BaAka, we have presented further evidence of a distinctive non-western gut microbiome, likely compatible with more ancestral human states. However, questions remain; although our data show that the BaAka microbiome is potentially active in fiber processing and energy harvesting, these results should be tested over a wider sample size, including different BaAka populations under dietary fluctuations and their Bantu neighbors at Dzanga. In this regard, the microbiome similarities seen between the BaAka and traditional African, South-American and Asian societies suggest that metabolic adaptations to phenolic and fiber-rich foods is not exclusive of the BaAka and that it may be a trait of traditional non-western groups. Also, the existence of an actual “ancestral” gut microbiome, characterizing these traditional non-western societies requires further exploration. This issue may be better understood by increasing efforts to reconstruct the vast uncharacterized diversity in these human populations and with comparative analyses that include wild nonhuman primates (56). Likewise, we urge the need to describe molecular interactions between diets, the gut microbiome and host immune markers in hunter-gatherers and industrialized societies. These approaches have the potential to unravel the benefits of increased gut bacterial diversity in traditional societies and may help us understand the forces that shaped modern microbiomes in the context of westernized disease patterns.

## Methods

### Study site, subjects and sample collection

Stool samples were collected from 27 BaAka. The group included 20 BaAka partially hired as gorilla trackers by the Primate Habituation Program (PHP) in DSPA and 7 female partners. The trackers alternate periods of time between the research camps, tracking the gorillas in the Park and their villages. Both trackers and their partners frequently do either day hunts from the villages or longer hunting trips to the Special Reserve. The samples were collected from June-August of 2011 upon consent from all individuals, and also as part of an effort to characterize their parasite loads. Approximately 1 g of feces was collected in 2-ml Eppendorf tubes containing RNAlater (Invitrogen, life technologies). Samples were kept at room temperature for a maximum of one month before transport to the Institute of Vertebrate Biology, Academy of Sciences of the Czech Republic, where they were kept at -20°C until they were shipped to the University of Illinois at Urbana-Champaign, USA where DNA was extracted. All work was approved according to rules and regulations from the Ministre de l’Education Nationale, de l’Alphabetisation, de l’Enseignement Superieur, et de la Recherche (Central African Republic) as well as the Institutional review board for the protection of human subjects from the University of Illinois at Urbana-Champaign, permit number 13045, September 4, 2014.

### Sample processing and DNA analyses

DNA was extracted from stool samples using the MoBio Ultraclean Soil kit (MoBio Laboratories Inc., Carlsbad, CA, USA). The V1-V3 region of the 16S rRNA gene was PCR amplified (20 cycles: at 94°C for 30s, at 48°C for 30s, and at 72°C for 2 min) using primers 27f (5′-AGAGTTTGATYMTGGCTCAG-3′, corresponding to nucleotides 8–27 of the *Escherichia coli* 16S rRNA gene) and 534r (5′-ATTACCGCGGCTGCTGGCA-3′, tagged with identifying barcodes, MID tag 1–50). The amplicons were multiplexed and pyrosequenced using 454 FLX-Titanium technology at the J. Craig Venter Institute (Rockville, MD, USA). After removing low quality sequences (<Q30, sequences shorter than 250unt, sequences with homopolymers longer than six nucleotides, and sequences containing ambiguous base calls or incorrect primer sequences) with custom-perl scripts, reads were processed using the online tool mothur, which included denoising, chimera check and denovo clustering of sequences sharing ≥ 97 % 16S rRNA sequence identity, into an operational taxonomic unit (OTU) (57).Taxonomic profiles of sequences was also conducted at genus, family, order, class and phylum level using the Ribosomal Database Project (RDP) Classifier (58) and the phylotype function within mothur. After sequence processing, OTUs detected fewer than five times across the entire data set and/or in fewer than 3 individuals were removed to avoid including probable sequence artifacts. For comparisons with the Hadza, Italians and Americans, we downloaded sequences deposited in MG-RAST, project ID 7058 (13), and from a subset of data available in the Human microbiome project (HMP) (http://www.hmpdacc.org/). 16S rRNA Sequences from Italians and the Hadza correspond to the V4 variable region obtained with 454 pyrosequencing. Sequences from US Americans, part of the HMP, correspond to the V1-V3 16SrRNA variable region, also obtain through 454 pyrosequencing. Thus, all sequence reads for the intergroup comparisons were analyzed using the closed reference OTU picking script against the greengenes database, as implemented in the QIIME pipeline (59). This pipeline was also used to calculate phylogenetic diversity as well as UniFrac distances. The PICRUSt (Phylogenetic investigation of communities by reconstruction of unobserved states) open source software (30) was used to predict the potential functional roles played by gut bacterial communities in the four groups. Sequence data from the BaAka have been deposited in MG-RAST, project ID XXXX.

### Metabolomic analyses

Extraction for polar and non-polar metabolites was performed separately. Metabolites were extracted with 1 ml of 70 % methanol and sonication in QSonica Microson XL2000 Ultrasonic Homogenizer (QSonica, LLC., CT, USA). Lysed cell pellets were subsequently fractionated at room temperature with 5 mL of 70% methanol, and chloroform, accompanied by centrifugation (10 min at maximum speed). One milliliter of each extract was evaporated under vacuum at -60°C and then dried extracts were derivatized (**See supplemental note for details**). The spectra of all chromatogram peaks were compared with electron impact mass spectrum libraries NIST08 (NIST, MD, USA), W8N08 (Palisade Corporation, NY, USA), and a custom-built library of 520 unique metabolites. All known artificial peaks were identified and removed. To allow comparison between samples, all data were normalized to the internal standard in each chromatogram and the sample dry weight (DW). The spectra of all chromatogram peaks were evaluated using the AMDIS 2.71 (NIST, Gaithersburg, MD, USA) program. Metabolite concentrations are reported as “(analyte concentration relative to hentriacontanoic acid) per gram dry weight” (relative concentration), i.e., as target compound peak area divided by the internal standard (IS) peak area (IS concentration is the same in all samples): N_i_ = X_i_ × X^-1^_IS_ × g wet w.

### Statistical analyses

All multivariate and community analyses were conducted using the *ca*, *vegan* and *labdsv* packages of R (60) on the relative abundance of each taxon (61-63). Indicator species analysis (64) was used to characterize the most prominent taxa in the gut microbiome of each group. Spearman correlations and Kruskal-Wallis tests were completed using the *psych* and *pgirmess* packages of R (65, 66). GC/MS spectra data from metabolomic analyses were transformed as follows: i) metabolites with >50% of missing data were removed from the set and ii) the relative abundance of a given metabolite was expressed relative to the sum of all the spectra obtained for a given sample.

## Acknowledgements

We would like to thank the government of the Central African Republic, namely the Ministre de l’Education Nationale, de l’Alphabetisation, de l’Enseignement Superieur, et de la Recherche for providing research permits to conduct our work in the Central African Republic; World Wildlife Fund and the administration of DSPA for granting research approval and for assistance with obtaining permits; and the Primate Habituation Programme for providing logistical support in the field. We thank Luis Barriero, George Perry, and Laure Segurel for important comments on this manuscript, as well all of our field assistants and trackers for their help in the field. This work was supported by NSF grant 0935347, Grant Agency of the Czech Republic (number 206/09/0927) and funds from the University of Minnesota College of Biological Sciences. This publication derives from the HPL-lab, Laboratory for Infectious Diseases Common to Humans and (non-Human) Primates, Czech Republic and was also co-financed by the European Social Fund and state budget of the Czech Republic. This work was supported in part by the University of Minnesota Supercomputing Institute.

The authors declare no conflict of interests.

